# Mapping immunological and host receptor binding determinants of SARS-CoV spike protein utilizing the Qubevirus platform

**DOI:** 10.1101/2023.07.27.550841

**Authors:** Carrie Sanders, Aristide Dzelamonyuy, Augustin Ntemafack, Nadia Alatoom, Godwin Nchinda, Millie Georgiadis, Alain Bopda Waffo

**Affiliations:** Center for Disease Control 1600 Clifton Rd NE, Atlanta, GA 30329; Department of Biochemistry and Molecular Biology, Indiana University School of Medicine, 635 Barnhill; Laboratory of Vaccinology and Biobanking, CIRCB BP 3077 Messa, Yaoundé, Cameroon

**Keywords:** Qubevirus, A1 minor coat protein, SARS-CoV, spike fragment, rhACE2, chimeric, anti-S antibody

## Abstract

The motifs involved in tropism and immunological interactions of SARS-CoV spike (S) protein were investigated utilizing the Qubevirus platform. We showed that separately, 14 overlapping peptide fragments representing the S protein (F1-14 of 100 residues each) could be inserted into the C-terminus of A1 on recombinant Qubevirus without affecting its viability. Additionally, recombinant phage expression resulted in the surface exposure of different engineered fragments in an accessible manner. The F6 from S_425-525_, was found to contain the binding determinant of the recombinant human angiotensin converting enzyme 2 (rhACE2), with the shortest active binding motif situated between residues S_437-492_. Upstream, another fragment, F7, containing an overlapping portion of F6 would not bind to rhACE2, confirming not just only that residues were linear but equally also the appropriate structural orientation of F6 upon the Qubevirus. The F6 (S_441-460_) and other inserts, including F7/F8 (S_601-620_) and F10 (S_781-800_), were demonstrated to contain important immunological determinants through recognition and binding of S protein specific (anti-S) antibodies. An engineered chimeric insert bearing the fusion of all three anti-S reactive epitopes, improved substantially the recognition and binding to their cognate antibodies. These results provide insights into humoral immune relevant epitopes and tropism characteristics of the S protein with implications for the development of subunit vaccines or other biologics against SARS-CoV.

**Significance:** Mapping epitopes within the receptor binding domains of viruses which are essential for viral tropism is critical for developing antiviral agents and subunit vaccines. In this study we have engineered the surface of Qubevirus to display a peptide library derived from the SARS-CoV S protein. In biopanning with S protein antibodies, we have identified three peptide fragments (EP1, EP2 and EP3) which reacted selectively with antibodies specific to the S protein. We demonstrated that all recombinant phage displayed peptide fragments both individually and as chimera exposed important immunological epitopes to their cognate antibodies. A peptide fragment F6 situated at S_425-525_, was found containing the binding determinant of the recombinant human angiotensin converting enzyme 2 (rhACE2), with the shortest active binding motif situated between residues S_437-492_. The platform is rapidly to identify epitopes and receptor binding sites within viral receptors found in target host cell. Thus, this platform holds great significance.

## Introduction

The severe acute respiratory syndrome (SARS) is caused by two major coronaviruses referred to as SARS-CoV and SARS-CoV-2 (1–5). The first SARS outbreak was in Guangdong province, in November 2002 (SARS-CoV) and the second in February 2020 (SARS-CoV-2) in Wuhan, China (6–10). The hallmark of both outbreaks was a rapid global spread of the disease thereby affecting several countries across the world (11–14). SARS-CoVs are enveloped, positive-sense RNA coronaviruses (CoVs) with a genome of about 30 kb in length (15,16). The genomes of both CoVs are similar in their organization and have several open reading frames encoding for the nuclear (N), membrane (M), envelop (E), and spike (S) proteins, respectively (17–20). The S protein is highly immunogenic and plays a crucial role in initiating viral infection through the recognition of its receptor, the angiotensin converting enzyme 2 (ACE2), expressed by the host cell (21–23). The S protein is common to SARS-CoV and SARS-CoV-2, with approximately 24.5% of non-conserved amino acid sequences (24, 25). Currently, although there is no approved vaccine against SARS-CoV, several effective vaccines have been approved against SARS-CoV-2. However, a continuous emergence of novel variants presents a formidable challenge not only in sustaining vaccine efficacy but also for developing new vaccines against both viruses. <u>In-silico</u> studies and computer prediction have mapped several domains of this multi-functional viral S protein that are involved in binding to ACE2 and in recognizing neutralizing anti-S antibodies (26,27). The multi-functionality of the S protein makes it druggable for prophylaxis and suitable for subunit vaccines development. In addition, known epitopes of the S protein could also be genetically engineered for diagnostic purposes.

The spike protein is one of the four major structural proteins of SARS-CoV which is characteristic of coronaviruses (28). Only 20-27% of amino acid homology was found while analyzing the S protein among coronaviruses (29). This difference in amino acid sequence is probably attributed to different features and functions. The S protein is a large glycoprotein incorporated into the viral envelope with two domains, S1 and S2, that exist as two non-covalently bonded subunits (30). The S1 is situated between residues 14 and 641 and consists of two subdomains, S1a and S1b (31), and is predicted to be responsible for the virus binding to its host cell receptor, ACE2 (32, 33). The binding motif of SARS-CoV to ACE2 happens via a putative binding fragment found on S1b which we have determined utilizing the RNA phage display system. The S2 is the transmembrane subunit and is made of two heptad repeat regions, HR1 and HR2 (34). The HRs facilitate viral and cellular membrane fusion in the fusogenic state (35). Receptor binding, as well as viral and host membrane fusion, are important steps in the virus cycle and pathogenesis. At the N- and C-termini of S protein are the signal and transmembrane peptides, respectively, which make S protein an attractive target for the development of antiviral agents. We have characterized the key peptide motifs using an evolutionary RNA phage display strategy.

During this study we have mapped immunological and host receptor-binding motifs of the SARS-CoV S protein following display upon the RNA coliphage display Qubevirus (Qβ) platform. This platform has recently been shown to expose several functional peptides without compromising the recombinant phage viability (36–40). Like CoVs, Qβ is a single stranded positive-sense RNA bacteriophage (41). Qβ belongs to the family of *Fiersviridae* and is small, being just 25 nm in diameter, with a 4.2 kb genome encoding for four proteins, including a replicase subunit (b), a major coat protein (Cp), a minor coat protein (A1 or MCP), and a maturation protein (A2 or MA2), respectively (42–46). The A1 was recently demonstrated by our group to be suitable for surface engineering for the exposition of recombinant epitopes (47–49, 36–40). Several expression cassettes within plasmids containing the full cDNA of the Qβ phage were constructed and used to insert and express over 100 amino acid long peptides at the C-terminus of the A1 protein.

Conventionally, the method of phage display panning immobilizes purified antibodies to plate or solid supports, to which the library is applied, and then the antibodies are extensively washed in a buffer solution containing detergent. To optimize the hybrid phage, a recovery form of the immobilized target needs to be undertaken with different elution conditions. Tight antibody-antigen (Ab-Ag) or protein-protein binding and attachment to the solid support can require a heated, low pH-value buffer treatment (50–54). This harsh treatment of sensitive Ab-Ag interaction can reduce the number of viable variants. We have succeeded in establishing an improved, optimized subtractive panning method to select and enrich antibody-specific antigens more efficiently without any elution (36).

During this study, a novel expression cassette was generated at the C-terminus of the A1 protein, which allows the insertion of 14 overlapping fragments of the S protein separately for recombinant phage production and target recognition analysis. Assuming that the phage expressed 12 copies of the recombinant A1 (36), the phage concentration of 10^4^ pfu/ml obtained for each fragment was used for panning against anti-S antibodies and for ELISA with a recombinant hACE2. One fragment motif was identified with binding activity to the hACE2, and three fragment motifs recognizing anti-S antibodies were also identified. A chimeric antigenic epitope consisting of the three fragment motifs displayed upon recombinant Qβ showed a high affinity for these same anti-S antibodies.

## Materials and Methods

### Phages, bacteria, plasmids, antibodies, and hrACE2

The Qubevirus or coliphage (Qβ) wild type was obtained from ATCC and the constructed QβAd2 with a deletion at the C-terminus of the A1 were maintained in the laboratory and used as controls for most experiments. For subcloning and plasmid maintenance, *E. coli* MC1016 was used. *E. coli* HB101 and DH5α were used to maintain recombinant plasmids containing the cDNA of Qβ and produce the first generation of phages from the corresponding plasmids. *E. coli* K12, Hfrh, and Q13 were indicator bacteria used to maintain, amplify, produce, and titer the phages. Plasmids pQβ8, pQβ7 (55), pBRT7Qβ, pQβAd2, and their recombinant derivatives were used for S protein fragment sequence insertion and recombinant phage display vector construction (56, 36–40). The antibodies were purchased from Sino Biological (Cat# 40150-MM08 and 40150-MM10) and ABclonal (Cat# RK04158) companies and a control human monoclonal antibody from SARS-CoV patients was provided by NIH (57). The recombinant human angiotensin-converting enzyme 2 (rhACE2) was purchased from Sino Biological (Cat# 10108-H08H).

### Construction of recombinant plasmid vectors for recombinant phages with various S protein fragments

Fusion PCR was used to generate each fragment insert containing, in frame with A1, its C-terminus, the linker peptide, and the fragment of S protein of SARS-CoV sequences, respectively. The first PCR with pQβ8 as template was done with the forward primer portion of the A1 sequence (1720–1767) containing the Bpu10I restriction enzyme site and a reverse primer containing the C-terminus of A1 (2282–2332) and the first 60 bp of each fragment sequence, respectively. The second PCR with the synthetic pUCS (S gene #) synthetic as template, was done with the forward primer containing the first 60 bp of each fragment and the reverse primer containing the last 50 bp of each fragment, the two natural stop codons of the A1 gene, a restriction enzyme site (PstI, EcoRV, NotI, NheI, or NdeI), the Shine-Dalgano gene, and the NsiI gene sequences, respectively. For each fragment, the final PCR was done with the purified products of the first and second PCR as templates and the Bpu10I-containing forward primer and the NsiI-containing reverse primer. For cloning, the final purified PCR product, and the vector pQβ8 were restricted with Bpu10I and NsiI and gel extracted. Additionally, the restricted vector was dephosphorylated and cleaned up with phenol, chloroform, and alcohol precipitation. The purified linearized vector and each fragment were separately ligated at 16 °C overnight as previously described (36, 37). For each S fragment, the total volume of ligation (20 µl) was used to transform 200 µl of competent *E. coli* MC1016 and plated. For each construction transformed, 5 to 10 clones were used to prepare DNA for screening. The DNA obtained was analyzed using the restriction enzyme added before the NsiI on the reverse primer. The confirmed DNA was subjected to Sanger sequencing to validate the S fragment inserted, the frame of the gene fusion, and the SD sequence for better expression. The recombinant plasmid with a positive S fragment was used for plasmid scaling and phage expression.

### Production of recombinant phages with various S protein fragments

For recombinant phage expression, production, and scaling, 20 ng of each recombinant DNA was used to transform *E. coli* HB101 or DH5α and plated on 2YT-agar as described elsewhere (36–39). For each recombinant plasmid with the corresponding fragment, two clone transformants were inoculated into 3 ml of 2YT supplemented with ampicillin and incubated at 37°C for 5 h while shaking at 150 rpm. Each initial culture was transferred to a liter of the same medium and incubate at 37°C overnight. The phages were extracted by PEG/NaCl precipitation as previously described (36–40). The phages obtained were titrated and used for large scale infection of *E. coli* K12 or Q13 at a multiplicity of infection (MOI) of 3. The phage titer was checked and used to determine the scale of amplification to the final titer of 10^12^-10^14^ pfu/ml. The amplified phages were precipitated as mentioned above and analyzed for the plaque’s morphology, concentration, and appropriate sequence.

### Panning selection of recombinant phages with S fragments against Anti-S antibodies

Recombinant phages with various S fragments obtained were used in this biopanning experiment. A high binding 96-well flat-bottom microplate was coated at 4°C overnight with 200 µl of anti-S antibody (2.5 µg/ml in coating buffer:15 mM Na_2_CO_3_ and 35 mM NaHCO_3_, pH 8.6). To block the empty surface of the wells, 100 µl of blocking buffer (2% BSA in PBST, pH 7.4) was added and incubated at room temperature for two hours. Excess unbound BSA was removed by washing twice with wash buffer (137 mM NaCl, 2.7 mM KCl, 8.3 mM Na_2_HPO^4^ 2H_2_O, 1.5 mM NaH_2_PO_4_ at pH 7.2 and 0.05% Triton X-A volume of 100 µl of recombinant phages (10^4^ pfu/ml) with S fragments preincubated in blocking buffer at 37°C for 1 h were then added to the wells and incubated at room temperature for 4 hrs. Unbound or loosely bound phage particles were extensively washed out with wash buffer 5 times. Bound phages were eluted by infection. A volume of 200 µl of log-phase *E. coli* Q13 were added to the wells and incubated at 37°C for 45 min. The bacterial culture from the experimental wells were then transferred into tubes and 100 µl titrated against *E. coli* Q13. The remaining aliquot was used as phage solution for the next round of panning against log-phase *E. coli* Q13. To ensure we have phages that bound tightly, more stringent washing conditions were used in subsequent rounds of panning. RT-PCR was used to characterize and validate the sequence of phages from each round of panning.

### ELISA and panning selection of recombinant phages with S fragments binding rhACE2 protein

#### ELISA of recombinant phages with S fragments

The binding effect of the S protein fragments to the host receptor rhACE2 was screened using an ELISA kit (CAYMAN, 205020) with slight modifications. Briefly, the recombinant phages were diluted in a coating buffer (NaHCO_3_) to a titer of10^2^ pfu/ml, and 100 µl was used to coat a 96-well plate at 4°C overnight. The coating buffer was removed, and wells were washed five times with the wash buffer, followed by blocking with 0.5% BSA in coating buffer for 1 h at room temperature. The buffer was discarded, and the wells were washed 5 times with the wash buffer. The rhACE2 was diluted at a concentration ranging from 0.078 to 2.5 µg/ml and 100 µl of each dilution was added into the well and incubated at room temperature for 1 h on an orbital shaker. The solution was discarded, and the wells were washed five times, 100 µl of antibody was added to the wells and incubated at room temperature for 1 h. The solution was discarded, the wells were washed 5 times and 175 µl of TMB was added. The plate was incubated for 30 min at room temperature and the reaction was stopped by adding 75 µl of the stop solution. The OD was recorded at 450 nm using a 96-well plate reader. Wells containing wild-type phage and 0.5% BSA were used as controls. The test was performed in triplicate at each concentration.

#### Panning selection of recombinant phages with S fragments

Biopanning was used to screen the recombinant phages against the rhACE2 as described elsewhere (36) with slight modification. Briefly, the wells were coated with 2.5 µg of rhACE2 as described above. The wells were washed three times with tris buffer saline containing 0.05% Tween-20 (TBST) and blocked with 0.5% BSA for 1 h at room temperature on an orbital shaker. The recombinant phage that showed binding to ACE2 in the ELISA assay was used, and 150 µl of the phage at a titer of 10^12^ was added to the wells. The plate was incubated in the same conditions as described above for 3 h. Thereafter, the wells were washed thrice with TBST and rinsed thrice with phage buffer followed by the addition of 150 µl of the host cell (*E. coli* Q13) at an OD_600_ 0.5-0.7. The plate was further incubated at 37°C on a shaker at 150 rpm for 1h and the phage titer of the first round was determined using plaque assay as described in our earlier publication (36).

### Confirmation of recombinant phage motifs with RT-PCR and dot blotting

#### RT-PCR

RNA of the recombinant phages was isolated using the QIAGEN kit. The cDNA was prepared and amplified by PCR as described elsewhere (40). The PCR product was purified from a 1% agarose gel and sequenced for confirmation of the inserted fragment within the recombinant phage genome.

#### Dot blotting analysis

Dot blotting analysis of the recombinant phages was performed as mentioned in our earlier study (40) except that phages at 10^13^ were used. After blocking the membrane, mouse IgG anti-S antibody at a dilution of 1:1000 was added and incubated at room temperature for 1h followed by washing 3x with TBST (TBS with 0.05% Tween-20). The bound anti-S antibody was further probed with anti-mouse HRP-conjugated at room temperature for 1h followed by washing 3x with TBST. The membrane was incubated with HRP substrate for 5 min at room temperature and exposed to chemiluminescence. The image was taken using Odyssey LI-COR Acquisition v1.2.0.72 software.

### Functional motif determination and structural analysis of recombinant phages with S fragments

Any fragment obtained having affinity with anti-S antibodies or rhACE2 was subjected to further analysis by residue deletion from the C-terminus and N-terminus, respectively. A maximum of 10 residues were removed from both ends of the fragment, and the binding function was accessed. For any loss of functionality, five residues were regained, and then one at a time until the full functionality was restored or improved. The corresponding motif was then sequenced, and the three-dimensional structure was determined. The sequence of the readthrough protein A1 and the engineered S determinant motifs were modeled with template-based modeling using the RaptorX web server, as reported elsewhere (37–40). All the obtained models were transformed to view the recombinant protein backbone and highlight the structure using the MolGro molecular viewer. The recombinant phage morphology was confirmed with cryo-electron microscopy (Cryo-EM), as described elsewhere (40).

## Results

### Construction of phage vectors for a SARS-CoV spike gene fragment library

We previously demonstrated that a consensus of 50 amino acids consisting of the membrane-proximal external-region (MPER) of the HIV-1 envelope gp41 could be built into the minor coat protein A1 for surface display upon recombinant Qβ phage (37). Using a similar approach in this study the C-terminus of the A1 protein was engineered to display a library of overlapping peptide fragments derived from the spike (S) protein of SARS-CoV (Figure 1L). Since the A1 is displayed on the surface of the phage particle, excessive modification might hamper recombinant phage production. As such, it was necessary for us to explore the tolerance of the Qβ genome for the insertion of longer DNA sequences. Since we had previously shown that a DNA fragment of 150 bp was stably inserted into the Qβ cDNA, the same procedure was thus sequentially used to generate recombinant phages bearing fragments of the S protein (36–40). During this process, restriction sequences of enzymes including Bpu10I and NsiI, were productively built into both the insert (each S fragment) and the plasmid vector (pQβ8). Likewise, plasmid vector expression cassettes were generated bearing an A1 gene modified in its C-terminus to display a library fragment of the S protein. As shown in Figure 2A, through sequential modification of the A1 genome, we have successfully increased from 150 bp to 300 bp the length of the DNA gene which can be fused with its C-terminus. The modified C-terminus of the 500 bp A1 genome was effectively fused with 300 bp and tolerance of the Qβ genome for such long inserted DNA (300 bp) was established. The overall impact of this process was the permanent modification of the A1 gene from 500 bp to 800 bp within the recombinant phage display vector generated for each fragment. Finally, a restriction site was built in between the A1 natural stop codons (TAG and TAA) and the NsiI cloning enzyme site prior to clones/plasmids analysis. A total of fourteen fragments of S protein were effectively fused to A1 separately and built into the phage cDNA. In Figure 2B and 2D, a restriction enzyme gel analysis is shown for each fragment with the expected length. Interestingly, the inter-regional section between A1 and the replicase genes was instead extended rather than reduced as previously reported (40). The recombinant plasmid library containing the various S fragments was successfully sequenced, analyzed, and shown to contain the desired designed frame and relevant features necessary for the expression of recombinant phages.

**Figure 1:**
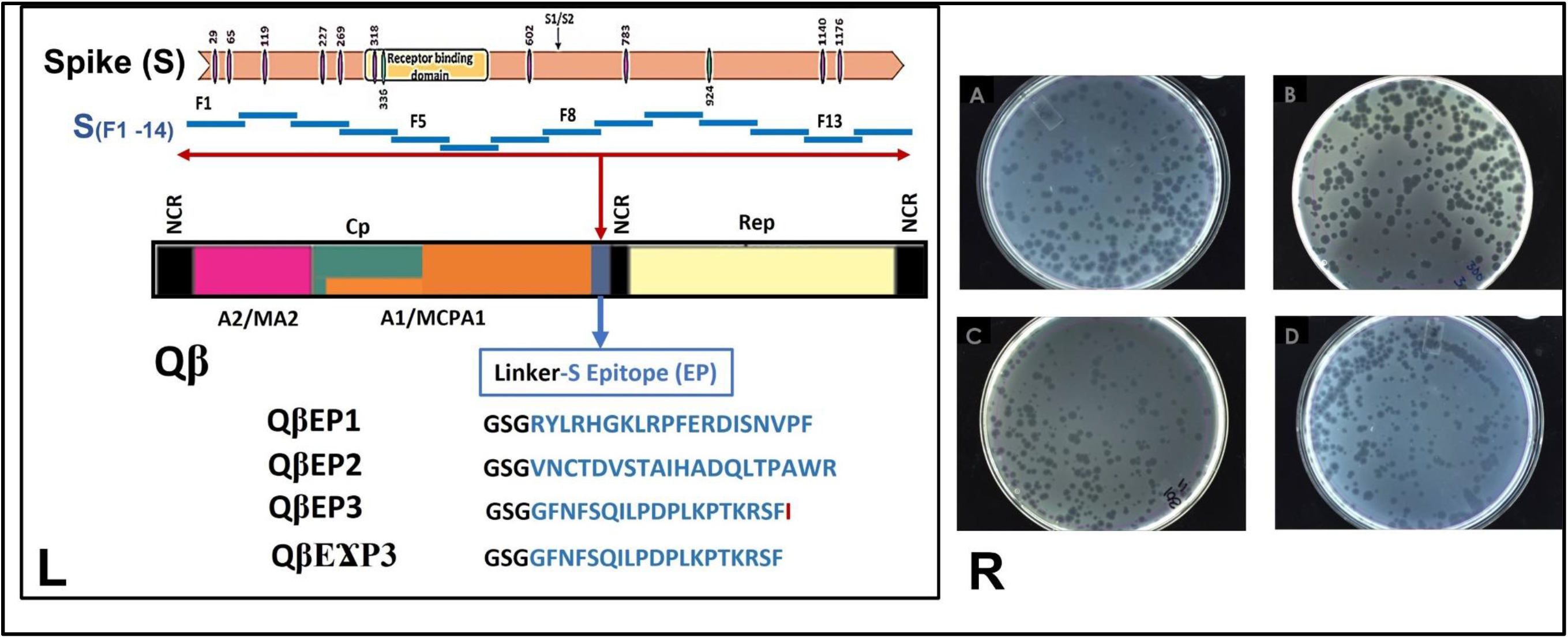
Epitope mapping scheme and plaque assay images. **(L):** Schematic representation of RNA coliphage Qβ insert of spike protein derived fragments from design to motif sequence selection. **UP**: Scheme of the known O-and N-glycosylation (purple and green bares) functional domains of the spike protein organization from the N-terminus signal sequence (F1), middle sequence with cleavage site (S1/S2) to the C-terminus (F14); **MIDDLE**: Phage Qβ genome organization of cDNA with non-coding region (NCR), maturation protein (A2/MA2), coat protein (Cp), read-through protein (A1/MCPA1) with the insertion cassette at the end (blue) for cloning, the replicase protein (Rep). **DOWN**: The residue motif of different epitopes (EP) obtained within the cassette on the Qβ phage; the phage with epitope 1 (QβEP1), the phage with epitope 2 (QβEP2), the phage with epitope 3 (QβEP3), the phage with epitope 3 with a deletion mutant of I (QβϪEP3). **(R):** Morphological image analysis of recombinant phages. In comparison to the wildtype (**Panel A**) the morphology of recombinant phages was analyzed on the lawn of <u>E. coli</u> Q13 host cell. **Panel B**: recombinant phage QβF6 harboring the fragment F6; **Panel C**: recombinant phage QβF8 containing the fragment F8; **Panel D**: recombinant phage QβF10 harboring the fragment F10. All results were observed after 6 hours of incubation at 37 °C.

**Figure 2:**
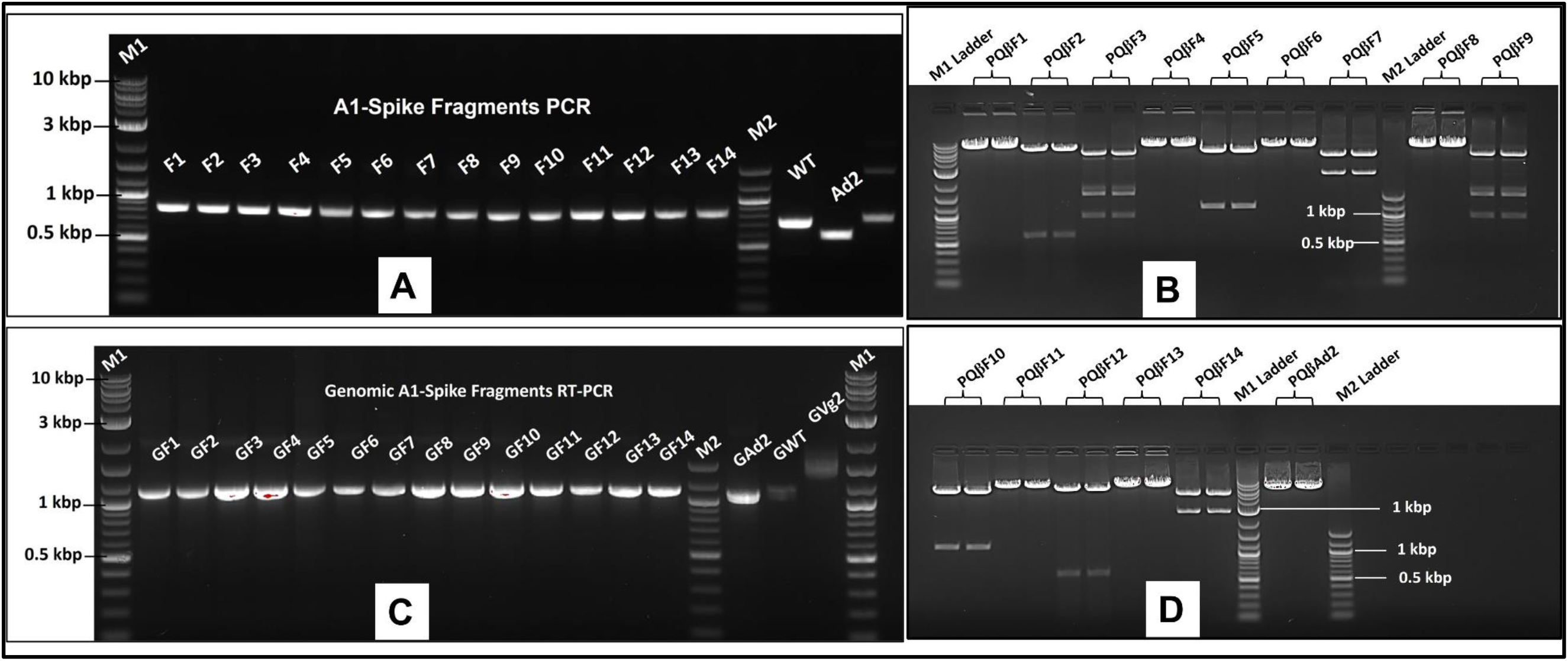
Agarose gel image analysis of the results of SARS-CoV S protein gene fragments. **(A)** Amplification: PCR products of 14 overlapping fragments amplification fusion PCR of a portion of A1 and of the S gene from F1 to F14 (∼800 bp); M2 and M1 are 10 kb and 100 bp ladder respectively. WT is the A1 gene amplified and Ad2 is the delete A1 gene amplified as control without the C-terminus 150 bp gene portion. **(B and D)**: Construction of recombinant phage vector for RNA phage display S fragments. Positive recombinant pQβF1, pQβF4, pQβF6, pQβF8, pQβF11, and pQβF13 plasmids are linearized with Not I respectively; positive recombinant pQβF2 and pQβF12 were digested into 700 bp and 7 kbp fragments with EcoRV respectively; positive recombinant pQβF3 and pQβF9 were digested into 1, 2.3, and 4.4 kbp with Nde I respectively; positive recombinant pQβF5 and pQβF10 were digested with Nhe I into 1 and 6.7 kbp fragments respectively; positive recombinant pQβF7 and pQβF14 were digested with Pst I into 3 and 4.7 kbp fragments respectively; M1 and M2 are 1 kbp and 100 bp DNA ladders respectively. The plasmid pQβAd2 is a negative control wildtype with A1 deletion linearized with Not I. **(C)** The RNA genotype size analysis for recombinant phages: From GF1 to GF14 are the genomic portion of the recombinant phages with S fragment 1 to 14 respectively, amplified by RT-PCR. GWT and GAd2 are the same genomic portion on the wild type and A1 deletion phages respectively, amplified by RT-PCR as negative controls. M1 and M2 are 1 kbp and 100 bp DNA ladders, respectively.

### Production of recombinant Qβ phage display library with various S derived fragments

To produce the first generation of recombinant Qβ phage, all plasmid vectors and constructed variants were transformed into *E. coli* DH5α or HB101, which are F^-^ bacteria lacking the pilus appendage for reinfection. Additionally, using F^-^ bacteria with plasmids with expression cassettes under the T7 promotor ensures exclusive usage of a high-fidelity DNA replication system that leads to plasmid transcription, resulting in a phage genome without premature evolutionary events within the phage expression system. Recombinant phages resulting separately from the expression of the recombinant plasmids with various S fragments were produced as plaques on lawns of their *E. coli* Q13 or K12 host (Figure 1R). Recombinant phage titers varying between 10^3^ - 10^5^ pfu/ml was obtained for various fragments, respectively (Table 1). The fragments 13 and 14 (F13 and F14) were poorly expressed and generated the lowest phage titer. The genome of each variant was analyzed by RT-PCR, agarose gel electrophoresis, and sequencing reactions. The results showed each variant containing the expected appropriate DNA fragment size (Figure 2C) when compared to the wild type and the control phage with a deleted A1. A fragment size of 1500 bp was formed consisting of the A1, S fragment, inter-regional section upstream of the replicase, and partial replicase genes, respectively. To confirm the presence of the corresponding S fragment gene within the plaques obtained each phage variant’s cDNA was sequenced. The results show that recombinant phages from each plaques corresponded specifically with their respective genes (genotype) fused in frame with the A1.

**TABLE 1:** Comparison of reactivity of different fragments of the SARS-CoV spike protein.

| Fragment | Position | Known as | Platform | Optimization | Plasmids | Phages | Titer<br>pfu/ml |
| --- | --- | --- | --- | --- | --- | --- | --- |
| F1 | S<sub>1-100</sub> | SP/NTD | -- | -- | pQβF1 | QβF1 | 10<sup>4</sup> |
| F2 | S<sub>85-185</sub> | NTD | -- | -- | pQβF2 | QβF2 | 10<sup>5</sup> |
| F3 | S<sub>170-270</sub> | NTD | -- | -- | pQβF3 | QβF3 | 10<sup>5</sup> |
| F4 | S<sub>255-355</sub> | NTD-RBD | No Binding | -- | pQβF4 | QβF4 | 10<sup>5</sup> |
| F5 | S<sub>340-440</sub> | RBD/RBM | No Binding | -- | pQβF5 | QβF5 | 10<sup>4</sup> |
| F6 | S<sub>425-525</sub> | RBM/RBD | Bind to<br>rhACE2 | 437-492 | pQβF6 | QβF6 | 10<sup>5</sup> |
| F6 | S<sub>425-525</sub> | RBM/RBD | IgG | 441-460 | pQβF6 | QβF6 | 10<sup>5</sup> |
| F7 | S<sub>510-610</sub> | S1 | IgG | 590-610 | pQβF7 | QβF7 | 10<sup>5</sup> |
| F8 | S<sub>595-695</sub> | S1 | IgG | 604-635 | pQβF8 | QβF8 | 10<sup>5</sup> |
| F9 | S<sub>680-780</sub> | S1/FP | -- | -- | pQβF9 | QβF9 | 10<sup>5</sup> |
| F10 | S<sub>765-865</sub> | FP/S2 | IgG | 785-800 | pQβF10 | QβF10 | 10<sup>4</sup> |
| F11 | S<sub>850-950</sub> | S2/HR1 | -- | -- | pQβF11 | QβF11 | 10<sup>5</sup> |
| F12 | S<sub>935-1035</sub> | HR1/S2 | -- | -- | pQβF12 | QβF12 | 10<sup>5</sup> |
| F13 | S<sub>1020-1140</sub> | S2 | -- | -- | pQβF13 | QβF13 | 10<sup>3</sup> |
| F14 | S<sub>1135-1255</sub> | HR2/TM/CP | -- | -- | pQβF14 | QβF14 | 10<sup>3</sup> |

### Selection of the S fragment(s) recognizing rhACE2 receptor and anti-S antibodies

We previously demonstrated a novel panning strategy whereby an FMDV major epitope’s synthetic library was selected using its specific cognate IgG monoclonal antibody immobilized on a plate resulting in a rapid fitness gain with just three rounds of biopanning (36). Separately during this study, recombinant human ACE2 (rhACE2), the natural receptor of SARS-CoV, and anti-S antibodies were immobilized as target prior to the selective biopanning procedure. All recombinant phages presenting various S fragments, respectively, were incubated either with rhACE2 or anti-S antibody (Table 1). After several washes to eliminate nonspecific binders, target bound recombinant phages (upon hACE2 or IgG) were amplified using a log phase culture of either *E. coli* Q13 or K12. Elution was achieved therefore by recombinant phage infection and the genomic characterization was followed by sequencing. Through this process the recombinant phages harboring an S fragment 6 (GF6) were identified which only bind specifically to rhACE2. On the other hand, other recombinant phages containing S fragments including 6, 7, 8, and 10 (GF6, GF7, GF8 and GF10), respectively were found to bind specifically to anti-S antibodies. The GF6 affinity to the anti-S antibodies was comparatively low requiring with more recombinant phages initially for selection. The selected recombinant phages bearing target specific S fragments were confirmed quantitatively by ELISA. The differential binding activity of recombinant phages to the anti-S antibodies and the rhACE2 is an indication that recombinant phages exposing the corresponding functional peptide sequences (phenotype) defined by the inserted gene in the phage genome.

### Determination of the S protein’s binding motif to the rhACE2 receptor

To determine the motif within S recognizing the rhACE2, the GF6 recombinant phages bearing the S fragment 6 were subjected to sequential deletion yielding mutants with 5, 3, and 2 residues from the N- and then C-termini. The mutant’s genes were synthesized by PCR and cloned into our optimized expression cassette. The recombinant plasmid was introduced into *E. coli* HB101 for expression and phage production. Each of the newly generated mutant recombinant phages were used for affinity analysis with rhACE2 as an agonist. A resultant motif with residues between 437-492 AA (S_437-492_) within GF6 (of S fragment 6) was identified as the smallest rhACE2-recognizing motif that retained the full selective binding activity using a quantitative ELISA. Any other deletion mutants bound weakly or almost lost the binding activity to rhACE2, indicating that the amino acids between positions 437-492 AA are essential in target host receptor recognition. The GF6 motif (S_437-492_) was subjected to panning using the immobilized rhACE2. Following elution by infection as previously described, recombinant phages, with a titer of 10^5^ pfu/ml was obtained with a round of amplification using *E. coli* Q13. The resultant recombinant phage was further subjected to quantitative ELISA using a two-fold serial dilution of rhACE2. The results showed a corresponding increase in absorbance (OD) of HRP conjugated anti-hACE2 antibodies with increasing concentration of rhACE2. A plateau was observed at a concentration of rhACE2 between 1.25 and 2.50 µg/ml (Figure 4B). The absorption curve was characteristic of the affinity between the SARS-CoV S receptor binding domain and the hACE2. Thus, our finding demonstrates the expected hACE2 binding motif within the S protein.

**Figure 4:**
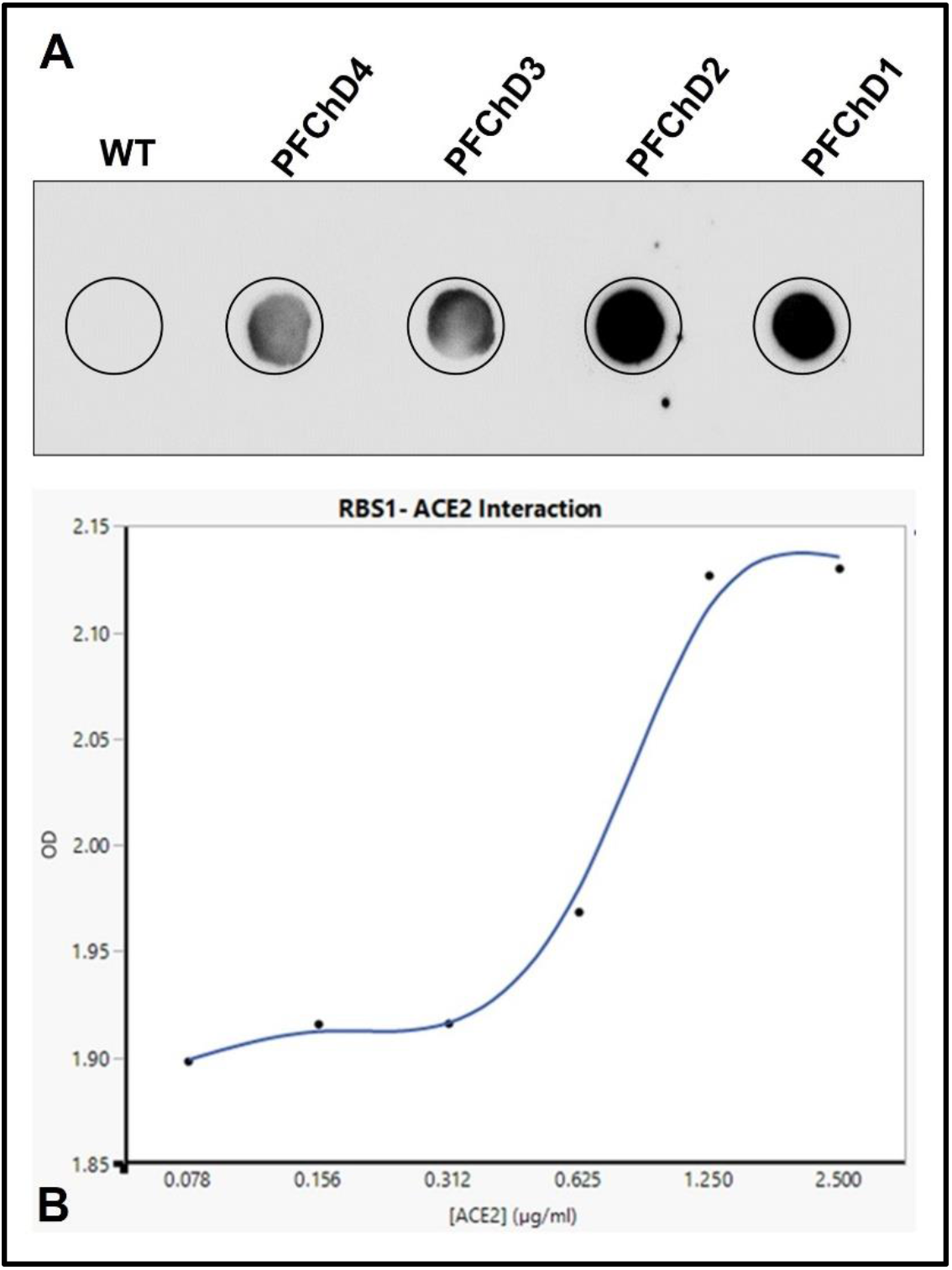
SARS-CoV S protein chimeric epitope reactivity analysis. **(A):** Dot blotting analysis of the recombinant purified phage harboring the chimeric epitope EPCh (chimeric of EP1, EP2, and EP3). WT is the control wild type; from PFChD4 to PFChD1 are 10^8^, 10^9^, 10^10^, and 10^11^ pfu/ml, respectively. **(B):** ELISA with recombinant phages QF6. The RBS1 - ACE2 interaction: the plot analysis of optical density (OD) of the horseradish peroxidase (HRP) product of anti-rhACE2 vs. increase of concentration of hACE2.

### Determination of the S protein motif recognizing anti-S antibodies

To determine the selective anti-S antibody reactive epitopes for the recombinant phages GF6, GF7, GF8, and GF10, a similar series of sequential N- or C-terminal deletion mutants of each fragment of the S region gene were generated by PCR and fused in frame with the A1 to reconstruct the corresponding recombinant plasmids, respectively. All these deletion mutants were performed sequentially from 10, 5, 3, and 2 residues from each end of the fragment and the affinity of the newly obtained recombinant phages to anti-S antibodies were analyzed, respectively. As a result, fragment 6 situated between residues 441-460 AA at position (S_441-460_) were found to retain a correspondingly increasing affinity for the anti-S antibodies. By deleting the C-terminus of F7 and the N-terminus of F8, the affinity to anti-S antibody was abolished, notably in quantitative ELISA, suggesting an overlapping region between both fragments. As for fragments 7 and 8 residues situated between 601-620 AA (S_601-620_) retained optimal affinity for anti-S antibodies while in fragment 10, it was instead residues 781-800 AA (S781-800). Sequentially, the residues starting from fragments 6, 7-8, and 10 were named epitope 1 (EP1), epitope 2(EP2), and epitope 3 (EP3), respectively. For all the epitopes obtained, deletion mutants of a single amino acid in the C-terminal dramatically weaken or almost completely abrogate the binding activity to the anti-S antibodies in contrast to similar deletions in the N terminal region. An example for EP3 is presented in Figure 1L, where a simple deletion mutant of isoleucine residue reduces the OD in ELISA by half. This indicates that the epitope is located at residues situated in the carboxyl terminal of EP1, EP2, and EP3, respectively, while the amino terminal is an extension of the platform used. The results of the fine epitope mapping are presented and summarized in Table 2.

**TABLE 2:** Comparison of residues position, sequence, and rank in affinity to anti-S antibodies of the different epitopes mapped of the SARS-CoV spike protein.

| Residue position | Amino acid sequence | Segment on the S protein | Rank in affinity |
| --- | --- | --- | --- |
| S<sub>441-460</sub> | RYLRHGKLRP FERDISNVPF | S1 | 1st |
| S<sub>601-620</sub> | VNCTDVSTAI HADQLTPAWR | S1 | 2nd |
| S<sub>781-800</sub> | GFNFSQILPD PLKPTKRSFI | S2 | 3rd |

### Analysis of recombinant phages bearing SARS-CoV S epitope motifs

The recombinant phages bearing mapped epitopes specific to anti-S antibody from the identified fragments of SARS-CoV S protein were subjected to further analysis using anti-S protein specific antibodies in ELISA and dot blotting. A three-dimensional (3D) computer simulation was used to analyze the fusion and exposition of the epitopes on A1. The 3D structural modeling indicated that each epitope is displayed without major impact on the A1 structure (color different from A1) on the C-terminal end of the A1 minor coat protein as shown (Figure 5LEFT). The results show that all three phages bearing epitope motifs (EP1, EP2, and EP3) interacted with the antibody with different binding affinity (Figure 3A). Their reactivity to the cognate antibody and the portion of the spike protein involved are depicted in Table 2, in comparison with other fragments and the wild type. By dot blotting visualization and quantitative ELISA analysis, EP1 showed the highest affinity to anti-S antibodies followed by EP2 and EP3, respectively. The results from dot blotting analysis were coincidental with those of ELISA (Figures 3A and 3B), indicating that the chimeric epitope recognized with a higher affinity the anti-S antibodies than the mapped single epitopes. This order of affinity for anti-S antibodies was substantially different among the corresponding fragments (F6, F7/8, and F10). This result demonstrates that a major anti-S specific epitope (EP1 of F6) is found in the S protein buried within the receptor binding domain and is successfully exposed using our recombinant phage platform. The result was in conformity with the 3D structural simulation of the recombinant A1.

**Figure 3:**
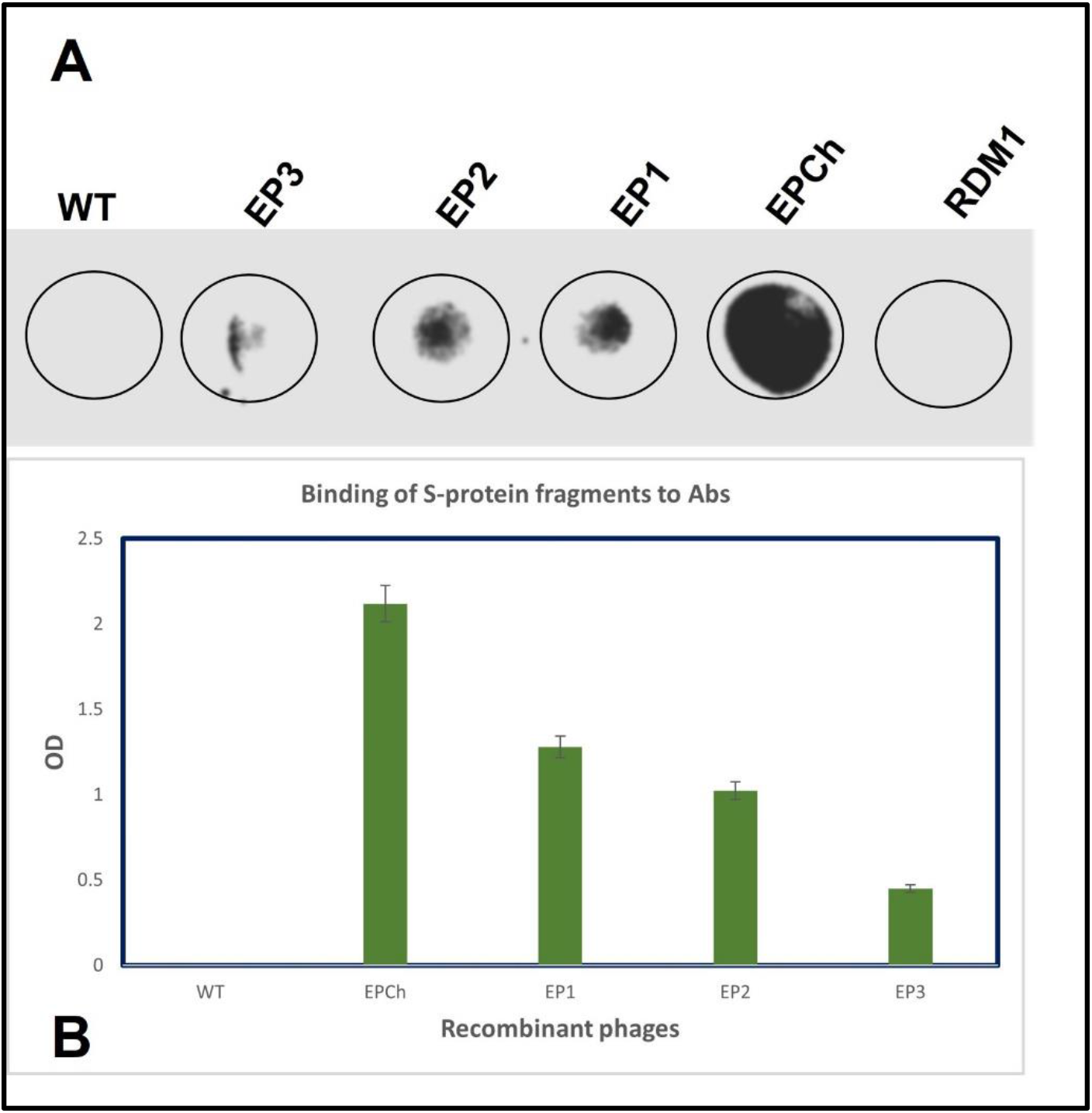
SARS-CoV S protein epitopes reactivity analysis. **(A)**: Dot blotting analysis of the recombinant purified phages harboring the epitope (EP) EP1 (F6: S_425-460_); EP2 (F7/F8: S_601-620_); EP3 (F10: S_781-800_); and EPCh (chimeric of EP1, EP2, and EP3). WT is the control wild type; RBM1 is the hACE2 receptor binding motif of SARS-CoV. The concentration of recombinant phages was 10^12^ pfu/ml, respectively. **(B):** ELISA Diagram analysis of S protein selected epitopes (EP). EP1 (F6: S_425-460_); EP2 (F7/F8: S_601-620_); EP3 (F10: S_781-800_); and EPCh (chimeric of EP1, EP2, and EP3). WT is the control wildtype. The concentration of recombinant phages was 10^5^ pfu/ml, respectively.

### Construction and analysis of recombinant phages bearing the chimeric epitope

A chimeric construct of all three epitopes (EPCh) was generated in sequential order, joined by linkers, and analyzed. EPCh which is the combination of the three other epitopes (EP1, EP2, EP3) described above showed strong binding activity to the antibody, followed by EP1, EP2, and EP3 in dot blotting analysis (Figure 4A). The same effect was observed in quantitative ELISA (Figure 4B), where the chimeric epitope (EPCh) showed the highest absorbance compared to the individual epitopes (EP1, EP2, EP3) as displayed on the surface of the phages. The binding activity of the chimeric epitope to the anti-S antibodies increased with the titer of the recombinant phage in dot blotting (Figure 5RIGHT). This result further confirms the reactivities of the epitopes mapped and also showed that a chimeric (all in one) may duplicate at least the binding reactivity of any of the single epitopes (EP1, EP2, or EP3) displayed upon recombinant phages. Moreso, the recombinant phages displayed epitopes showed no effect on the viability, structure, morphology, and stability when compared to the wild type.

**Figure 5:**
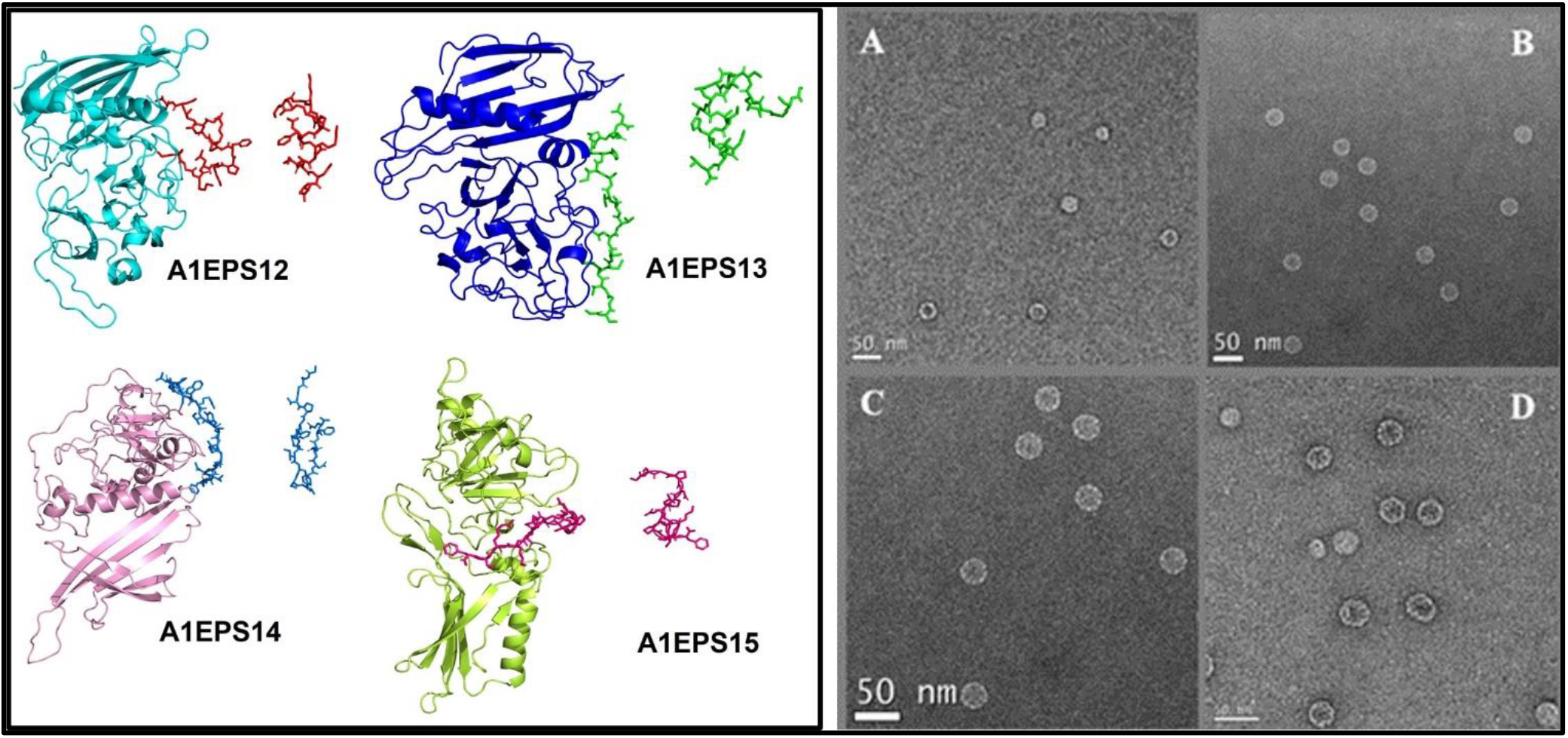
**(LEFT)**: Three-dimensional representation of structure prediction of the recombinant minor coat protein A1. In (A1EP1), QβEP1; (A1EP2), QβEP2; (A1EP3), QβEP3; (A1ϪEP3), QβϪEP3. For each, the fragment depicted is short in a different color from A1, which is large. **(RIGHT)**: Cryo-electron microscope image. (A) Qβ wild type; (B) QβF7; (C) QβF8; (D) QβF10.

## Discussion

Since the 2003 SARS and 2019 COVID19 disease outbreaks, coronaviruses have been isolated, characterized, and sequenced. However, little is known about their molecular tropism, antigenicity, and mechanism of pathogenesis. Additionally, with the knowledge generated from previous studies about other coronaviruses demonstrate that the spike protein plays a vital role in mediating viral cell entry through binding to its receptor and contributes to the induction of neutralizing antibodies. Computer prediction and simulation studies have been done to determine the potential S protein receptor binding domain and epitopes. Receptor binding motif are residues of the virus structural S protein which are functional determinants involved in initiating the tropism and thus, are druggable. Epitopes within the S protein induce antibody production and cellular immunity against viruses. These epitopes can be targeted for antiviral drugs, subunit vaccines, and point-of-care diagnostic kit development. In this study, we have generated recombinant Qβ phages displaying a library of peptide fragments from the SARS-CoV S protein. In combination with its commercially available stable cellular receptor (rhACE2), we identified a peptide sequence hereby referred to as fragment 6 (GF6) located between residues 437-492 AA within the SARS-CoV S protein selectively binding its cognate host cell receptor. When the recombinant Qβ-S phage library was assessed with target specific anti-S antibodies and monoclonal antibodies from SARS patients (kindly provided by NIH), three reactive epitopes of SARS-CoV S including EP1, EP2, and EP3 were identified. These epitopes were all located in the S protein between peptide residues AA 441-460, 601-620, and 781-800, respectively.

Since the receptor binding motif is essential for recognition and adsorption to the target cells, we hypothesized that our identified GF6 with selective binding to rhACE2 should be located within the receptor binding domain (RBD) of S protein. In the recombinant Qβ phage library a GF7 fragment downstream of the GF6 with overlapping peptides would not bind to the SARS-CoV receptor rhACE2. This probably suggests a conformational orientation of the binding motif within the fragment. Similar to GF6, its optimized derivative motif showed proportional rhACE2 concentration dependent binding activity, thereby demonstrating the selective interaction with its cognate receptor. When 100 recombinant phage particles were immobilized and exposed to this motif, binding saturation was achieved with 1.2 µg/ml of rhACE2 protein. Deletion mutants generated by targeting the binding motif of GF6 beyond the amino acid at positions 437-492, bound weakly or almost lost the binding reactivity with rhACE2.Thus confirming that the receptor binding motif is located within the SARS-CoV S protein. Previously, Babcock et al. (58) reported that the S-derived fragment consisting of amino acid residues between 270 and 510 efficiently bound ACE2 with a higher degree than the complete receptor binding domain of S1. Our results provide important information for drug design against the SARS disease. The A1 gene was engineered to accommodate up to 300 bp of additional sequences, which was above the usual fragment insertion size, indicating that the Qβ phage can tolerate longer genes (up to 300 bp for now) and still produce viable visible phage plaques. The production of visible plaques resulting from recombinant phages harboring S fragments, is a demonstration of the tolerance of the minor coat protein A1 to manipulation for subsequent insertion of additional nucleotides. In the A1-modified recombinant phages, additional insertions neither disturbed their functionality nor viability. However, some minor differences were observed in plaque morphology between various inserted fragments. The varying plaque morphology was nevertheless predominated by plaques with medium size in diameter (2 mm). Normally plaque morphology is determined by the insert length and structure, however, since all the S-derived fragments were similar in length, the observed differences could be ascribed to the quasispecies nature of the RNA virus population. Thus, a quasispecies occurrence is a consensus of sequences within the same variant. The A1 protein fused to 100 additional amino acids, can still participate in the virions formation (expression and assembly) and confer to the recombinant phage its infectivity, although with a reduced phage titer. Recombinant Qβ phages presenting this binding motif can be used immediately for a competition study with the pseudo-virus within our core facility. Historical precedent exists which demonstrated safe administration of phages to humans during the pre-antibiotic era for therapeutic purposes to patients (59). Therefore, a potential exists for the possible use of phage displaying agonists such as the receptor binding motif of SARS in patients to compete with SARS-CoV for binding to its host receptor hACE2 in clinical applications.

The antigenicity of proteins is determined by specific epitopes playing a key role in defining their reactivity with the target host immune system. During antibody-antigen reaction, unique epitopes located in specific portions of the antigen are vital in driving selective reactivity within their cognate antibody. This definition implies that the epitope and its cognate antibody must interact by a lock and key mechanism. In this wise, antigen may contain several epitopes corresponding to different antibodies. Two S protein-specific antibodies alongside with a monoclonal antibody from SARS patients were used for this study. SARS-CoV S protein-derived fragments 6, 7, 8, and 10 (GF6, GF7, GF8, and GF10) showed selective reactivity with the S protein-specific antibodies. Whereas GF6 bound weakly, the others showed significant higher selective reactivity during panning and in ELISA assays. Additionally, the receptor-binding motif also bound weakly to the S protein-specific antibodies. Mapping the S fragments using sequential deletion mutants identified an epitope EP1 within GF6 with a strong binding activity to S specific antibodies, followed by two other epitopes EP2 and EP3. This indicates that the reactivity may solely not depend only on the primary structure of the displayed S fragment. These results suggest that EP1 is probably unfavorably buried within the receptor binding motif, which probably in effect also hinders reactivity to its cognate antibodies. Normally, EP1, EP2, and EP3 are linearly situated within the S protein sequence, and after fusion to A1 on the Qβ platform, they retain their requisite conformation thereby efficiently recognizing S protein-specific antibodies. For each epitope, the binding reactivity to the S protein-specific antibodies was different, which probably suggests a complementarity between them. For all the S fragment motifs identified, the carboxyl terminal was essential for epitope reactivity to its cognate antibody. Thus, deletion mutants of just a single amino acid at the carboxyl terminal led to a dramatic weakening or complete abrogation of reactivity to S protein-specific antibodies. The deletion of an isoleucine (I) for example at the C-terminus of EP3 resulted in a reduction by half of its OD during quantitative ELISA assay. This single deletion mutant of EP3 probably results in a loss in its conformational structure, which is necessary for its stability during intramolecular interaction while reacting with S protein-specific antibodies which in effect also, dampen its reactivity. Hua et al. (60) have mapped two linear epitopes recognizing monoclonal antibodies D3C5 and D3D1, corresponding to 447-458 and 789-799 amino acids of the S protein of SARS-CoV, respectively. EP1, EP2, and EP3 identified in this study further highlight the function of S during SARS-CoV pathogenicity.

An S protein derived chimeric epitope (EPCh) engineered to include all three optimized epitopes was constructed, expressed, and used for binding in antibody reactivity assays as previously described. EPCh is a discontinuous epitope since it is formed by residues that are not contiguous in the amino acid sequence of the S protein but are brought together through genetic fusion with a linker as a spacer. EPCh bound comparatively stronger than EP1, EP2, and EP3 to S protein-specific antibodies in both quantitative ELISA and dot blotting. EPCh showed proportional binding activity for all cognate antibodies with increasing recombinant phage titers. This augmented reactivity in functional assays is certainly due to the fact that recombinant phages harboring chimeric epitopes present an increased number of epitopes, which as a consequence also increases the avidity. Computer simulation analysis have shown that EP1 and EP2 are in the spherical head of the S protein, while EP3 is in the stem portion. We showed recently that A1 occupies the 12 corners of the Qβ phage icosahedral structure. Such positioning mimics and exposes the S protein derived functional motifs on the surface of the recombinant phage particle. The Qβ phage has successfully served as a platform for the display of all the motifs identified during this study and provided a linkage between residues displayed and their appropriate genes; thus, this connection between the genotype and the phenotype can be easily manipulated to obtain desirable nanotools.

In conclusion, we have identified through recombinant Qβ phage mimicry the fine S protein-specific antibody epitopes to be situated at residues AA 441-460, 601-620, and 781-800 respectively, which are different from those previously known. These epitopes were confirmed by monoclonal antibodies from SARS patients (kindly provided by NIH). Additionally, a fragment containing amino acid residues between 437-492 is described, which selectively binds to rhACE2. Thus, our results provide useful biochemical information for an in-depth understanding of SARS-CoV S protein and its exploitation for the development of both prophylaxis and therapeutics against SARS-CoV infection.

## AUTHOR CONTRIBUTIONS

A. B. W., A. D., and C. S. generated the library of S fragments and variants of the ORF used for phage display selection assays; A. B. W. and M. G. conducted the structural and Cryo-EM analyses; C. S., N. A., A.N., and A. D. conducted initial phage purification and analyses. A. N., N. A., and G. N. conducted dot blotting and ELISA analyses; A.B.W., A.N., and A.D. conducted the phage vector plasmid construction and expression; A.B.W., A. D., and G.N. conducted phage display simulation and restriction assays; A.B.W. and G.N. wrote the original manuscript draft; all authors participated in finalization of the manuscript and approval of its final full text and figures.

## ACKNOWLEDGEMENTS

The phage library construction and screening were done within the Indiana University School of Medicine (IUSM) Chemical Genomics Core Facility. The cryo-EM images performed by the iCEM core of IUSM. Our gratitude to the Indiana University Precision Health Initiative (PHI) for the start-up fund.

